# Compartmentalized CRISPR Reactions (CCR) for High-Throughput Screening of Guide RNA Potency and Specificity

**DOI:** 10.1101/2024.05.07.592954

**Authors:** Tinku Supakar, Ashley Herring-Nicholas, Eric A. Josephs

## Abstract

CRISPR ribonucleoproteins (RNPs) use a variable segment in their guide RNA (gRNA) called a spacer to determine the DNA sequence at which the effector protein will exhibit nuclease activity and generate target-specific genetic mutations. However, nuclease activity with different gRNAs can vary considerably, in a spacer sequence-dependent manner that can be difficult to predict. While computational tools are helpful in predicting a CRISPR effector’s activity and/or potential for off-target mutagenesis with different gRNAs, individual gRNAs must still be validated in vitro prior to their use. Here, we present compartmentalized CRISPR reactions (CCR) for screening large numbers of spacer/target/off-target combinations simultaneously in vitro for both CRISPR effector activity and specificity, by confining the complete CRISPR reaction of gRNA transcription, RNP formation, and CRISPR target cleavage within individual water-in-oil microemulsions. With CCR, large numbers of the candidate gRNAs (output by computational design tools) can be immediately validated in parallel, and we show that CCR can be used to screen hundreds of thousands of extended gRNA (x-gRNAs) variants that can completely block cleavage at off-target sequences while maintaining high levels of on-target activity. We expect CCR can help to streamline the gRNA generation and validation processes for applications in biological and biomedical research.

## MAIN TEXT

The Clustered Regularly Interspaced Short Palindromic Repeats (CRISPR) and CRISPR-associated protein 9 (Cas9) system (Figure 1) has emerged as a powerful tool for targeted gene editing [1, 2], allowing precise and efficient modifications of specific nucleic acid sequences within a genome. The process relies on a cofactor of the Cas9 protein known as a guide RNA (gRNAs) that is designed to be complementary to targeted DNA sequences in a modular segment of the RNA called their spacer [3, 4]. Stable base-pairing between the spacer sequence and the target sequence activates the Cas9’s nuclease domains to cleave the DNA [3-7]. If Cas9 cleavage occurs within a cell, cellular double-strand break repair (DSB) mechanisms can introduce permanent genetic changes into its genome at the site of the break. Therefore, by simply changing the sequence of the gRNA’s spacer, Cas9 can be used to introduce specific mutations into a gene of interest [8, 9, 10, 11] which often involves insertions or deletions during mutagenic DSB repair [8, 9] that can cause a frameshift knockout of the gene, making it a very useful tool for biological studies and for gene therapies [1,2].

**Figure 1.**
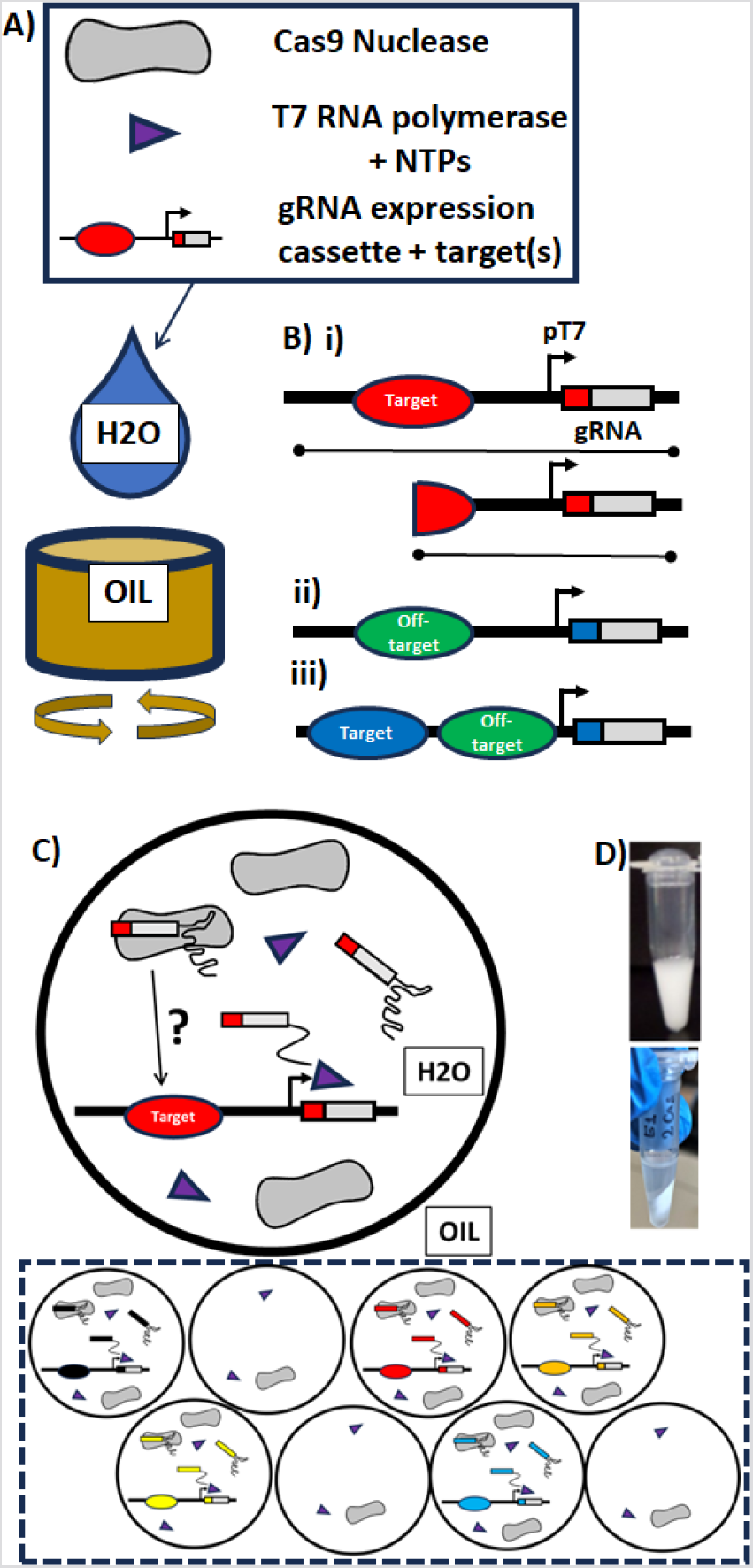
A) Compartmentalized CRISPR Reactions (CCR) are performed in water-in-oil emulsions containing high concentrations of Cas9 nucleases and T7 RNA polymerases with nucleotide triphosphates (NTPs) for RNA transcription, and low concentrations of DNA substrates so that individual DNA molecules are expected to be isolated within a single emulsion. B) The DNA substrates are designed to contain a module for T7 RNA polymerase based expression of a gRNA and (i) a target for Cas9 with that gRNA, so that activity of the Cas9 within that emulsion that with specific gRNA can be determined from the length of the resulting DNA substrates with that spacer sequence (red); (ii) a potential off-target site (green) for a gRNA of interest (blue); or (iii) a combination of both, where DNA molecules predominantly exhibiting cleavage patterns at the spacer-distal site would imply a gRNA with significantly higher activity on-target than at spacer-proximal off-target sites. C) (above) Within each emulsion, unique gRNAs are transcribed from isolated DNA molecules, where they can associate with a Cas9 nuclease then either demonstrate or fail to demonstrate cleavage activity at a (off- or on-) target site of interest. (below) If a library of different DNA molecules with different gRNA/target/off-target combinations (different colours) are used when generating the water-in-oil emulsions, large numbers of gRNA/target/off-target combinations can be evaluated simultaneously. D) (left) Photograph of a water-in-oil emulsions in a microcentrifuge tube; (right) after the emulsions have been broken after the the CCR reaction via centrifugation for subsequent analysis.

However, the molecular process of target recognition and cleavage by Cas9 is complex and dynamic: the Cas9 enzyme recognizes a short motif known as a PAM [7] immediately next to the target sequence in the DNA and opens the double helix, allowing the gRNA’s spacer sequence to displace the non-targeted strand nucleotide by nucleotide [4], forming a structure known as an R-loop [12, 13]. The formation of a stable R-loop structure drives a conformational change in the Cas9 that positions the nucleases appropriately to generate the DNA DSB. As a result of this dynamic process, predicting the ability of Cas9 to recognize and degrade different sequences is more difficult than, say, design of PCR primers which is driven largely by the thermodynamics of base-pairing [14, 15]; instead, this process, known as strand invasion by the gRNA, is believed to be a kinetic, non-equilibrium process, and the ability of Cas9 to recognize and introduce DSBs into DNA can vary significantly based on multiple properties of the gRNA’s spacer sequence [16-26] and its ability to form a stable R-loop after strand invasion [3, 4, 5, 12, 13]. Cas9 RNPs will exhibit little or no nuclease activity for many gRNAs [16-26]. Significant efforts have gone into predicting the activity of Cas9 RNPs with different gRNAs [16-26], and machine learning models [23, 24, 25, 26] have emerged as useful tools for determining gRNA candidates to knockout a gene of interest, however these models often do not have excellent predictive power. Predictions of nuclease activity by these models have shown a correlation coefficient of only around 0.4 with experimental results, and they depend strongly on the experimental system to which they are compared [32].

The nuclease domains of Cas9 can also be activated If, during strand invasion, the gRNA is able to stably bypass mis-paired nucleotides at off-target sequences that are similar but not quite identical to the intended target sequence. Therefore, off-target effects are also considered when determining a gRNA for a target gene of interest to prevent unwanted mutagenesis [16,18, 25-31]. In practice, trade-offs must often be made between the predicted on-target activity and the potential for off-target activities [16, 18, 25-32]. For CRISPR-based knockout of a gene of interest, computational tools will typically (i) enumerate all possible target sequences within the exon sequences of that gene by the presence of a requisite PAM; (ii) predict activity using one of many machine learning models; (iii) identify the possible off-target sequences by sequence similarity to the target and for each gRNA predict nuclease activity at those sites; then (iv) output the results [32].

It can therefore be difficult to translate the output result of computational tools, as different models for on-target, off-target, and DSB repair outcomes can contradict one another [32], and multiple gRNAs must be tested or validated in vitro by hand prior to use in order to efficiently generate the desired mutations in cells [33, 34, 35]. Conducting this validation for multiple gRNAs within cells one at a time can be cumbersome [36-39], and often in vitro screening [34] is performed to assess nuclease activity using purified RNPs as a standard course. This too requires transcribing, purifying, and combining gRNAs individually with Cas9. Assessing large numbers of spacer/target/off-target combinations presents many practical difficulties [40, 41] for the purpose of determining most effective and most precise gRNA for a given application.

Here, we present a streamlined approach, inspired by the concept of compartmentalized self-replication (CSR) [41-50] to simultaneously assess CRISPR nuclease activity and specificity in vitro for multiple gRNA in parallel, in a process that we call a Compartmentalized CRISPR Reaction (CCR) screen (Figure 1). During a CCR screen, libraries of short (<500 bp), specially designed DNA molecules (Figure 1B) are individually confined within micron-sized water droplets in water-in-oil emulsions along with high concentrations of T7 RNA polymerase and Cas9 nuclease. The emulsification process involves slowly adding the aqueous phase in a dropwise manner to a stirring mixture of mineral oil and surfactants. The DNA molecules contain the coding sequence for a gRNA downstream of a promoter for T7 RNA polymerase, as well as a targeted DNA sequence (or off-target) on the opposite end of the DNA molecule. Under buffer conditions we have identified where both T7 RNA polymerase and Cas9 were found to be active (Supporting Information, Figure S1), the gRNAs encoded on the isolated DNA molecules are transcribed and can associate with the Cas9 nucleases to form the active Cas9 RNP, which can then interact with the target sequence found on the same molecule within the emulsions (Figure 1C). After breaking the emulsions (Figure 1D), removing protein, and determining the length of the DNA molecules and the sequence of the spacer on the segment coding for the gRNA via run-off next generation sequencing, we can determine the activity of many gRNAs simultaneously, in a single one-pot reaction. This allow us, for example, here to (i) test the activities of all the gRNAs in the first exon of a gene of interest experimentally, (ii) evaluate activities at the likely off-target sequences for a gRNA at the same time, and (iii) screen hundreds of thousands of gRNAs variants for those with greater specificities and activities than a standard gRNA for a target of interest (see below).

Approximately 90% of the water-in-oil emulsions droplets were ∼850 nm in diameter on average (+/-218 nm, standard deviation (stdev.)) measured via dynamic light scattering (DLS), with ∼10% being larger (∼4.2 um in diameter, +/-1.042 um stdev) (Supporting Information, Figure S2). A back-of-the-envelope calculation therefore suggested that there were 3.5 x 10^14 water-in-oil emulsions per mL [40], and that adding approximately 0.1 - 1 femtomole of the DNA (∼600 million to ∼6 billion molecules) per mL would ensure a ratio of 1 DNA molecules for every 100 – 1000 emulsions, even within the large droplets if present, which we hypothesized would essentially ensure that each DNA molecule would be isolated within its own emulsion while still allowing us to test a large number of gRNA/target/off-targets independently and simultaneously (Figure 1C). To test whether these emulsions would successfully isolate the independent gRNA generation, RNP formation, and nuclease events, we designed a pair of DNA molecules each containing a gRNA expression cassette, its target on the distal end of the DNA molecule, and the target for the paired gRNA positioned proximally between its target and the promotor (Figure 2). This way, if each of the DNA molecules were isolated in their own emulsion, we might expect to see DNA with cuts only on at the distal target sites, while if both species of the DNA pair were mixed (each in the same emulsion), we might expect to see DNA that were cut at the proximal target site (Figures 2A and 2B).

**Figure 2.**
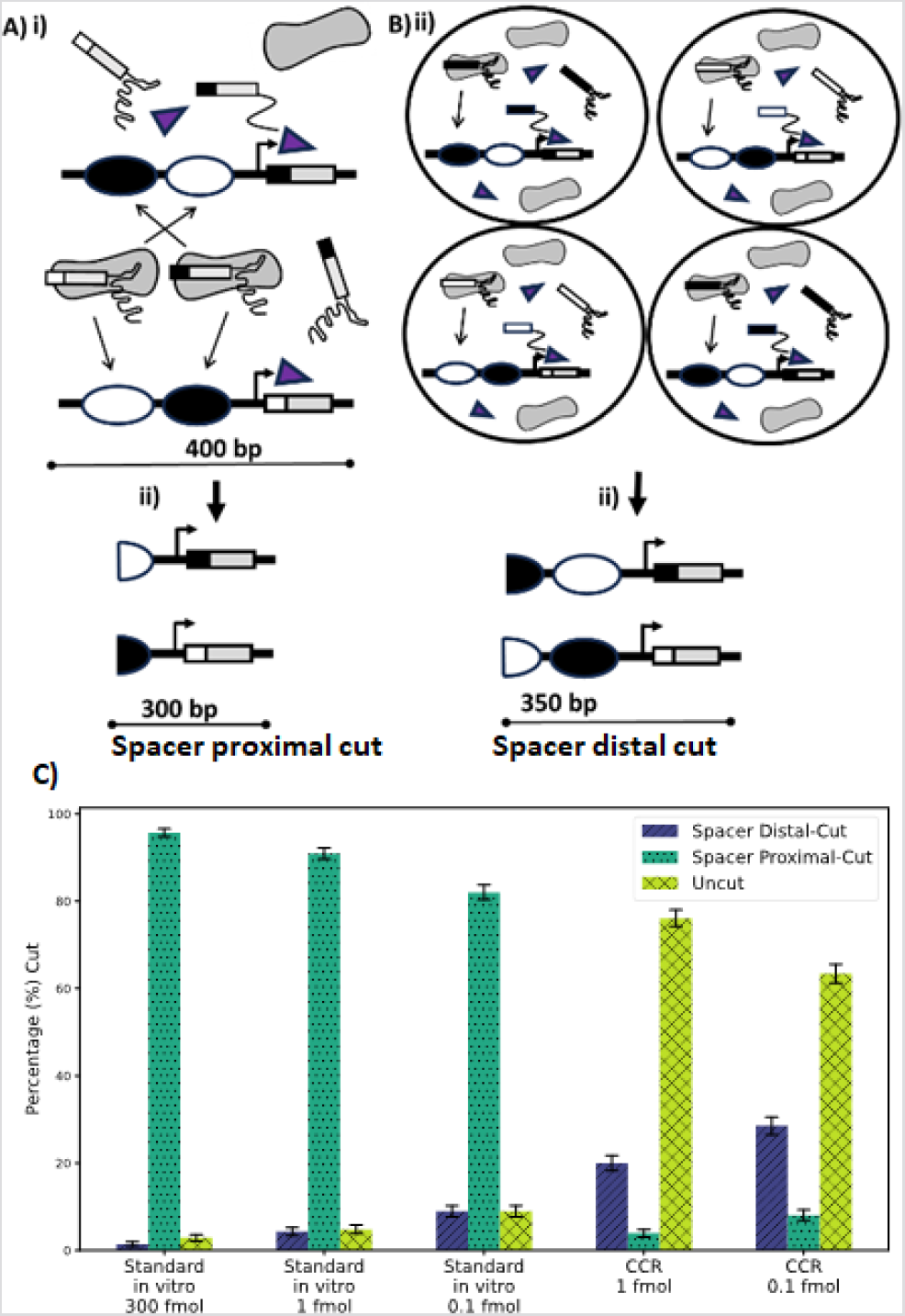
Confirmation that CCRs are performed with individual DNA molecules isolated within separate water-in-oil emulsions. A) Two DNA molecules are designed: each with a target (black or white) specific to the gRNA encoded on that molecule located on the distal end of the DNA (black or white, respectively) and the paired target for the other located proximal to the spacer (white or black, respectively). Under conditions where the two molecules were mixed, one would expect to see all of the DNA molecules cut at the proximal sites (300 bp), as the paired gRNAs could cause Cas9 to introduce DSBs at those sites, regardless of whether there were DSBs at the distal sites. B) Under conditions where the DNA molecules were isolates in emulsions, after CCR reactions one would expect mostly DNA molecules with DSBs at the distal site (∼350bp). C) Under standard (with no emulsions) conditions where the molecules were introduced to both Cas9 and T7 RNA polymerase, after the reaction most of the DNA molecules are determined to be cleaved at the spacer proximal site. However, those DNA molecules reacted within water-in-oil emulsions were found to be significantly less likely to be cleaved at the spacer-proximal site, and the vast majority of those that had been cleaved were cleaved at the spacer-proximal target, strongly implying that individual DNA molecules were isolated during the CCR reaction (See also supporting information, Figure S3).

We performed this in vitro reaction either under standard in vitro reaction conditions (with no emulsions) (Figure 2A) or within the emulsions (Figure 2B) and determined the size of the resulting DNA molecules after an 8-hour reaction period (Figure 2C). Those DNA molecules under standard conditions were essentially all cut at the proximal target site, as both gRNAs were generated together and CRISPR RNPs could easy access both respective targets on either molecule. In the emulsions, the overall fraction of cleaved molecules decreased, but those that were cleaved were mostly cleaved at the distal sites (Figure 2C and Supporting information, Figure S3). This implied that most of the DNA molecules were isolated in individual emulsions that greatly reduced the potential for cross-reaction between RNPs with gRNAs generated from one DNA and the targets positioned on another DNA molecule.

An advantage of CCR is elimination of the need of cloning, and the libraries of individual DNA molecules needed for CCR can be synthesized inexpensively. These libraries can also be easily scaled to many members of the library as a pooled set of oligonucleotides, each containing both a unique target candidate and its paired gRNA candidate, that can be amplified using PCR to generate the double-stranded DNA molecules needed for CCR. Considering this, we then tested the utility of the CCR approach for larger screens by evaluating the on-target activity of all 18 possible spacers (Figure 3A) [16-25] that are capable of targeting sequences within the 126 bp exon 1 of the human gene EMX1 (Chr2:72933825-72933951 in the hg38 assembly). Of those, one gRNA (gRNA 16) has been historically well-characterized for both on- and off-target activities [51, 52]. CRISPOR, a computational tool for gRNA design, by default reports the efficiency scores or prediction of activity from two common machine learning approaches known as Doench ‘16 (sgRNA Designer) [16] and Moreno-Mateos (CrisprScan) [17], and it also reports prediction scores for 8 other previously published methods [18-25]. As can be seen (Figure 3A), the different prediction methods are generally inconsistent with one another, with no one gRNA having high predicted efficacy scores across all methods, a situation which can complicate interpretation of these outputs and decision-making regarding which gRNAs are most likely to result in an active RNP. According to trend observed following the CCR screen of all 18 gRNAs simultaneously (Figure 1C and 3B), among the top performing RNPs, one includes the well-characterized gRNA (gRNA 16) which we have confirmed is active. Furthermore, the in vitro (using purified RNP complexes reacting individually with DNA molecules) validation of two gRNAs with demonstrated higher activity during CCR screen (gRNAs 12 and 10) revealed that they indeed had higher on-target activity compared to gRNA 16 (Figure 3B inset). These results show that CCR offers a straightforward approach for high-throughput screening and ranking of on-target activity, starting from a pooled oligonucleotide library.

**Figure 3.**
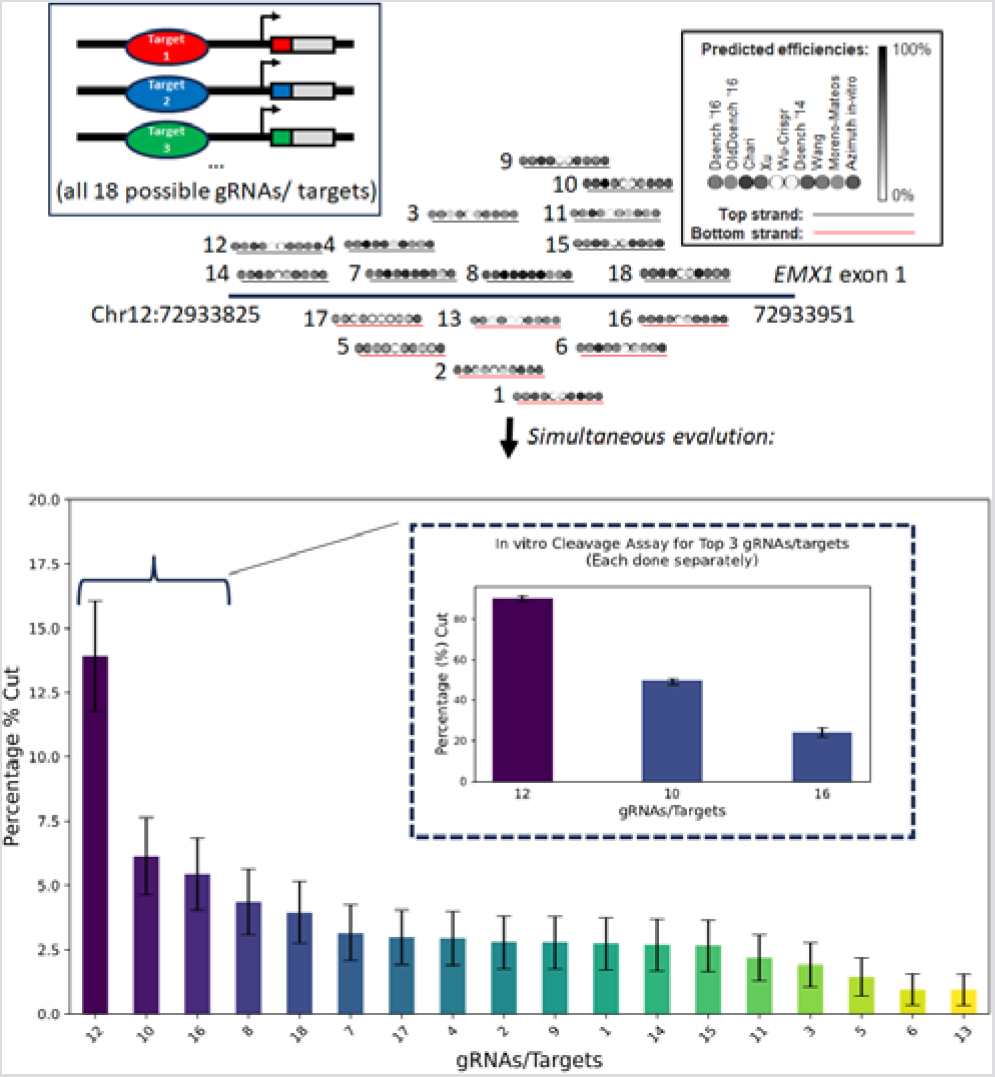
CCR to screen all gRNAs for a target gene in a one-pot reaction. A) The overlapping positions of all 18 possible gRNAs for exon 1 of EMX1, with computationally predicted activity for Cas9 at those sequences from 10 different models[16-25] to predict gRNA activity. As can be seen, there is little agreement across models, which can make interpretation difficult. B) Observed cleavage rates for all 18 gRNAs reveals a gRNA known to be active (gRNA 16) to be among the best performing. Error bars are binomial confidence (95%) based on the number of next-generation sequencing reads for each gRNA/target DNA molecule. (inset) In vitro validation using purified RNPs shows the trend observed with CCR recapitulated the results from more labour-intensive screening for RNP activity with different gRNAs one at a time. Error bars are std across 3 replicates (See also supporting information, Figure S4).

In some circumstances, it may be desired or advantageous to simultaneously screen both on- and off-target propensities (Figure 4A), because even if a gRNA is highly active, it is not useful if it is also active at the potential off-target sites. To do so, we could use a DNA molecule similar in design to those used in Figure 2, with the on-target site located distal to the gRNA expression region and the off-target located proximally. This way, after screening, the identity of the spacer with the resulting length of the DNA molecule can be determined: if the DNA molecule cut only at the distal target, that is evidence that the RNP with that gRNA is active and specific, since the off-target site remains uncut; but if it is cut at the proximal off-target, it indicates that the RNP with that gRNA is not specific enough, even if we do not know the activity on-target (if the molecule is uncut that is evidence of poor activity of the RNP with that gRNA). Following this design we also screened four of the gRNAs from CRISPOR [32] with low predicated off-target activities, as well as gRNA 16 which is associated with several off-target sites with known off-target activity, totalling 18 predicted off-target sites (Figure 4B). The CCR method re-capitulated the relative off-target activity at four known off-targets for gRNA 16 and mirrored its in vitro activity at these sites [52] (Figure 4B inset), further demonstrated that CCR could be used to screen a large array of off-targets as well. The off-target activity for the other gRNAs largely followed trends predicted by computational tools CFD and MIT off-target scores, but surprisingly gRNA 2, which computationally is not predicted to have a high amount of activity at off-target sites, exhibited the highest levels of activity during CCR at those off-target sites (Figure 4B). This highlights the importance of experimentally validating the computational predictions regarding gRNAs prior to use, which CCR makes significantly easier.

**Figure 4.**
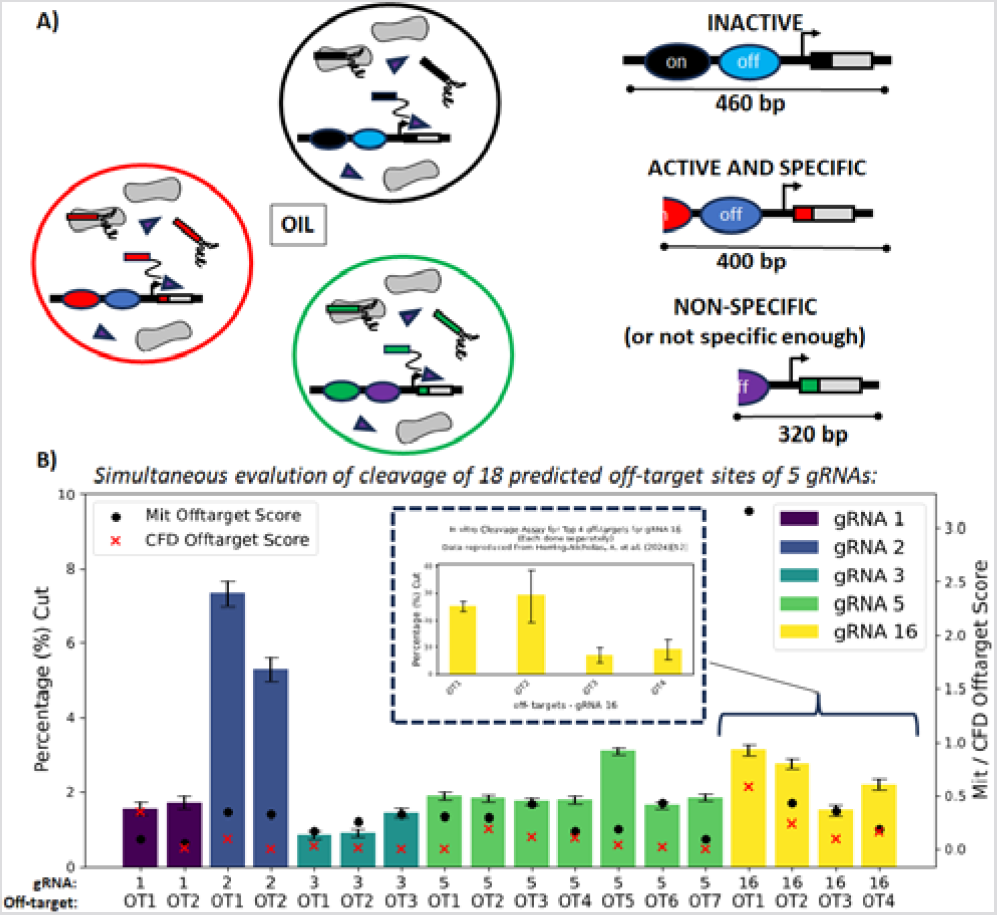
Screening different gRNAs for off-target activity simultaneously in parallel using CCR. A) Cleavage rates at the off-target sites of the different gRNAs, 5 predicted to have very low off-target activity (1, 2, 3, and 5) and one with known off-targets (gRNA16), with predicted off-target scores (right, arrows and dots). Error bars are binomial confidence (95%) based on the number of next-generation sequencing reads for each gRNA/target DNA molecule. (inset) Off-target activity for gRNA 16 using CCR follows that pattern observed when activity at those sites is validated one-at-a-time using purified RNPs.

We then tested the effectiveness of CCR in identifying highly active and specific gRNA variants by screening a large library of gRNAs with 8 nucleotide 5’-extensions (x-gRNAs). As previously, reported [51,52], gRNAs with short (8-12) nt extensions to the 5’-end immediately next to their spacer sequence are capable of eliminating the off-target activity of an RNP with that x-gRNA while still maintaining its on-target activity as the original gRNA. However, the task of identifying the specific sequence of the 5’-nt’s can be challenging, as many extensions either have no effect on specificity or also attenuate on-target activity. To screen the x-gRNAs library, we created a DNA library encompassing all possible 5’-extensions of 8 nt’s (65,536 possible sequences) for EMX1 gRNA 16, which exhibits well documented activity at several known off-target sites within the human genome, as re-capitulated through CCR method, on a DNA molecule which had its target on the distal end and one each of the four most active off-target sites at the proximal end (Figure 5A): a total of 262,144 separate gRNA/target/off-target combinations within a single CCR reaction. Following CCR screen, the top five x-gRNAs were identified from the most common x-gRNA spacer sequences with spacer-distal cleavage patterns (Figures 5A, S5, and S6). Their efficacies were further validated in vitro using purified RNPs and it was found that they were able to completely block activity at the off-target site, while maintaining on-target activity comparable to that of a high-specificity engineered variant of Cas9 known as eCas9 (Figure 5B). Therefore, CCR can be an effective tool to screen large numbers for gRNA variants with special attributes as well.

**Figure 5.**
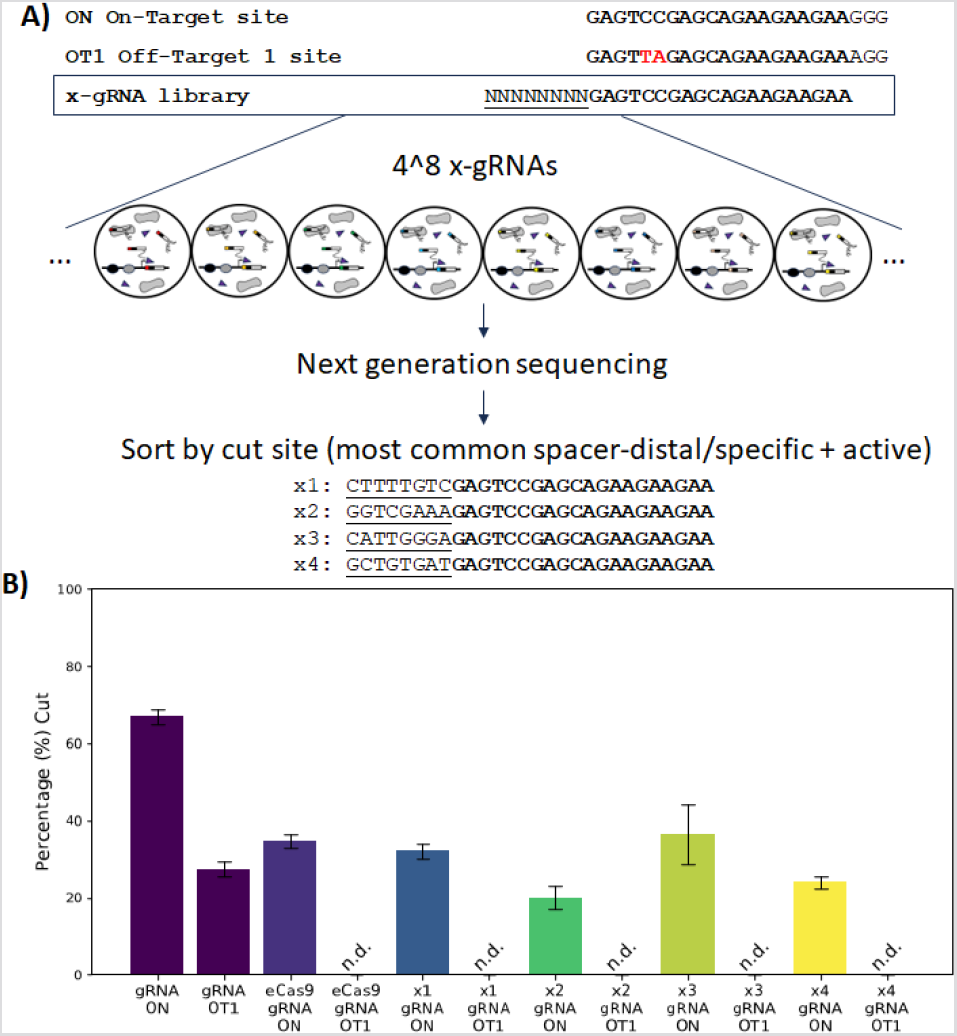
CCR to screen hundreds of thousands of gRNA variants for high activity and specificity. A) gRNAs with 5’ extensions of 8 nucleotides (x-gRNAs) are known to be capable of increasing the specificity of the gRNA, although identifying the precise sequence of the x-gRNA can be difficult. With CCR, all 4^8 x-gRNA sequences can be evaluated simultaneously. B) Validated in vitro one at a time, x-gRNAs identified by having the most common specific and active (spacer-distal) cleavage pattern all exhibited on-target (ON) activities comparable to an engineered high-specificity Cas9 (eCas9) while not having any detectable nuclease activity at their off-target site (OT1).

In conclusion, water-in-oil emulsions technique have been useful for handling large sets of unique DNA molecules in parallel, such as in library preparation for next-generation sequencing (NGS) [41, 42, 43] and in directed evolution studies [44, 46, 48], and here we show that the complete CRISPR reaction (gRNA transcription, RNP formation, and target cleavage) can be reconstituted within individual emulsions to screen large libraries of gRNAs/targets/off-targets in a streamlined and experimentally simplified manner. CCR represents powerful yet accessible method to easily screen many candidate gRNAs (provided from the outputs of computational tools for gRNA design) for both activity and specificity, and as a way of rapidly identifying highly active and specific gRNA variants from large sets of gRNA/target/off-target combinations. We expect CCR can be applied to screen large sets ofgRNAs for other CRISPR effectors as well, particularly considering that the available computational tools for gRNA design are currently advanced for only a select few effectors, while new effectors with diverse properties are increasingly discovered [53].

## Acknowledgements

The authors with to acknowledge financial support from NIH National Institute of General Medical Sciences (NIGMS) (R35GM133483), NIH National Institute of Biomedical Imaging and Bioengineering (NIBIB) (R21EB033595), as well as NSF (award #32027738) which provided support for T.S and DoD (Contract #W911QY2220006) which provided support for AHN. This work was performed in part at the Joint School of Nanoscience and Nanoengineering, a member of the Southeastern Nanotechnology Infrastructure Corridor (SENIC) and National Nanotechnology Coordinated Infrastructure (NNCI), which is supported by the NSF (Grant ECCS-1542174).

## References

1. Ran, F. A., Hsu, P. D., Wright, J., Agarwala, V., Scott, D. A., & Zhang, F. (2013). Genome engineering using the CRISPR-Cas9 system. Nature Protocols, 8(11), 2281–2308.

2. Hsu, P. D., Lander, E. S., & Zhang, F. (2014). Development and applications of CRISPR-Cas9 for genome engineering. Cell, 157(6), 1262–1278.

3. Jiang, F., & Doudna, J. A. (2017). CRISPR-Cas9 Structures and Mechanisms. Annual Reviews of Biophysics, 46, 505–529.

4. Sternberg, S. H., Redding, S., Jinek, M., Greene, E. C., & Doudna, J. A. (2014). DNA interrogation by the CRISPR RNA-guided endonuclease Cas9. Nature, 7490, 62–67.

5. Jinek, M., Jiang, F., Taylor, D. W., Sternberg, S. H., Kaya, E., Ma, E., Anders, C., Hauer, M., Zhou, K., Lin, S., & Kaplan, M. (2014). Structures of Cas9 endonucleases reveal RNA-mediated conformational activation. Science, 343(6176), 1247997.

6. Nishimasu, H., Ran, F. A., Hsu, P. D., Konermann, S., Shehata, S. I., Dohmae, N., Ishitani, R., Zhang, F., & Nureki, O. (2014). Crystal structure of Cas9 in complex with guide RNA and target DNA. Cell, 156(5), 935–949.

7. Anders, C., Niewoehner, O., Duerst, A., & Jinek, M. (2014). Structural basis of PAM-dependent target DNA recognition by the Cas9 endonuclease. Nature, 513(7519), 569–573.

8. Kosicki, M., Tomberg, K., & Bradley, A. (2018). Repair of double-strand breaks induced by CRISPR–Cas9 leads to large deletions and complex rearrangements. Nature biotechnology, 36(8), 765–771.

9. Brinkman, E. K., Chen, T., de Haas, M., Holland, H. A., Akhtar, W., & van Steensel, B. (2018). Kinetics and fidelity of the repair of Cas9-induced double-strand DNA breaks. Molecular cell, 70(5), 801–813.

10. Allen, F., Crepaldi, L., Alsinet, C., Strong, A. J., Kleshchevnikov, V., De Angeli, P., Páleníková, P., Khodak, A., Kiselev, V., Kosicki, M., & Bassett, A. R. (2019). Predicting the mutations generated by repair of Cas9-induced double-strand breaks. Nature biotechnology, 37(1), 64–72.

11. Gagnon, J. A., Valen, E., Thyme, S. B., Huang, P., Ahkmetova, L., Pauli, A., Montague, T. G., Zimmerman, S., Richter, C., & Schier, A. F. (2014). Efficient mutagenesis by Cas9 protein-mediated oligonucleotide insertion and large-scale assessment of single-guide RNAs. PloS one, 9(5), e98186.

12. Jiang, F., Taylor, D. W., Chen, J. S., Kornfeld, J. E., Zhou, K., Thompson, A. J., Nogales, E., & Doudna, J. A. (2016). Structures of a CRISPR-Cas9 R-loop complex primed for DNA cleavage. Science, 351(6275), 867–871.

13. Mullally, G., Van Aelst, K., Naqvi, M. M., Diffin, F. M., Karvelis, T., Gasiunas, G., Siksnys, V., & Szczelkun, M. D. (2020). 5′ modifications to CRISPR–Cas9 gRNA can change the dynamics and size of R-loops and inhibit DNA cleavage. Nucleic acids research, 48(12), 6811–6823.

14. Rychlik, W. (1993). Selection of primers for polymerase chain reaction. PCR protocols: current methods and applications, 31–40.

15. Mann, T., Humbert, R., Dorschner, M., Stamatoyannopoulos, J., & Noble, W. S. (2009). A thermodynamic approach to PCR primer design. Nucleic acids research, 37(13), e95–e95.

16. Doench, J. G., Fusi, N., Sullender, M., Hegde, M., Vaimberg, E. W., Donovan, K. F., Smith, I., Tothova, Z., Wilen, C., Orchard, R., & Virgin, H. W. (2016). Optimized sgRNA design to maximize activity and minimize off-target effects of CRISPR-Cas9. Nature biotechnology, 34(2), 184–191

17. Moreno-Mateos, M. A., Vejnar, C. E., Beaudoin, J. D., Fernandez, J. P., Mis, E. K., Khokha, M. K., & Giraldez, A. J. (2015). CRISPRscan: designing highly efficient sgRNAs for CRISPR-Cas9 targeting in vivo. Nature methods, 12(10), 982–988.

18. Stemmer, M., Thumberger, T., del Sol Keyer, M., Wittbrodt, J., & Mateo, J. L. (2015). CCTop: an intuitive, flexible and reliable CRISPR/Cas9 target prediction tool. PloS one, 10(4), e0124633.

19. Chen, W., McKenna, A., Schreiber, J., Haeussler, M., Yin, Y., Agarwal, V., Noble, W. S., & Shendure, J. (2019). Massively parallel profiling and predictive modeling of the outcomes of CRISPR/Cas9-mediated double-strand break repair. Nucleic acids research, 47(15), 7989–8003.

20. Xu, H., Xiao, T., Chen, C. H., Li, W., Meyer, C. A., Wu, Q., & Liu, X. S. (2015). Sequence determinants of improved CRISPR sgRNA design. Genome Research, 25(8), 1147–1157.

21. Wong, N., Liu, W., & Wang, X. (2015). WU-CRISPR: characteristics of functional guide RNAs for the CRISPR/Cas9 system. Genome Biology, 16, 1–8.

22. Doench, J. G., Hartenian, E., Graham, D. B., Tothova, Z., Hegde, M., Smith, I., Sullender, M., Ebert, B. L., Xavier, R. J., & Root, D. E. (2014). Rational design of highly active sgRNAs for CRISPR-Cas9–mediated gene inactivation. Nature biotechnology, 32(12), 1262–1267.

23. Wang, D., Zhang, C., Wang, B., Li, B., Wang, Q., Liu, D., Wang, H., Zhou, Y., Shi, L., Lan, F., & Wang, Y. (2019). Optimized CRISPR guide RNA design for two high-fidelity Cas9 variants by deep learning. Nature communications, 10(1), 4284.

24. Fusi, N., Smith, I., Doench, J., & Listgarten, J. (2015). In silico predictive modeling of CRISPR/Cas9 guide efficiency. BioRxiv, 021568.

25. Chari, R., Yeo, N. C., Chavez, A., & Church, G. M. (2017). sgRNA Scorer 2.0: a species-independent model to predict CRISPR/Cas9 activity. ACS synthetic biology, 6(5), 902–904.

26. Chuai, G., Ma, H., Yan, J., Chen, M., Hong, N., Xue, D., & Liu, Q. (2018). DeepCRISPR: optimized CRISPR guide RNA design by deep learning. Genome Biology, 19, 1–18.

27. Listgarten, J., Weinstein, M., Kleinstiver, B.P., Sousa, A.A., Joung, J.K., Crawford, J., Gao, K., Hoang, L., Elibol, M., Doench, J.G., & Fusi, N. (2018). Prediction of off-target activities for the end-to-end design of CRISPR guide RNAs. Nature biomedical engineering, 2(1), 38–47.

28. Bae, S., Park, J., & Kim, J. S. (2014). Cas-OFFinder: a fast and versatile algorithm that searches for potential off-target sites of Cas9 RNA-guided endonucleases. Bioinformatics, 30(10), 1473–1475.

29. Hsu, P.D., Scott, D.A., Weinstein, J.A., Ran, F.A., Konermann, S., Agarwala, V., Li, Y., Fine, E.J., Wu, X., Shalem, O., & Cradick, T.J. (2013). DNA targeting specificity of RNA-guided Cas9 nucleases. Nature Biotechnology, 31(9), 827–832.

30. Pliatsika, V., & Rigoutsos, I. (2015). “Off-Spotter”: very fast and exhaustive enumeration of genomic lookalikes for designing CRISPR/Cas guide RNAs. Biology direct, 10, 1–10.

31. Xie, S., Shen, B., Zhang, C., Huang, X., & Zhang, Y. (2014). sgRNAcas9: a software package for designing CRISPR sgRNA and evaluating potential off-target cleavage sites. PloS one, 9(6), e100448.

32. Haeussler, M., Schönig, K., Eckert, H., Eschstruth, A., Mianné, J., Renaud, J.B., Schneider-Maunoury, S., Shkumatava, A., Teboul, L., Kent, J., & Joly, J.S. (2016). Evaluation of off-target and on-target scoring algorithms and integration into the guide RNA selection tool CRISPOR. Genome Biology, 17, 1–12.

33. Sentmanat, M. F., Peters, S. T., Florian, C. P., Connelly, J. P., & Pruett-Miller, S. M. (2018). A survey of validation strategies for CRISPR-Cas9 editing. Scientific reports, 8(1), 888.

34. Tsai, S. Q., Nguyen, N. T., Malagon-Lopez, J., Topkar, V. V., Aryee, M. J., & Joung, J. K. (2017). CIRCLE-seq: a highly sensitive in vitro screen for genome-wide CRISPR–Cas9 nuclease off-targets. Nature methods, 14(6), 607–614.

35. Arndell, T., Sharma, N., Langridge, P., Baumann, U., Watson-Haigh, N. S., & Whitford, R. (2019). gRNA validation for wheat genome editing with the CRISPR-Cas9 system. BMC biotechnology, 19, 1–12.

36. Zhou, Y., Zhu, S., Cai, C., Yuan, P., Li, C., Huang, Y., & Wei, W. (2014). High-throughput screening of a CRISPR/Cas9 library for functional genomics in human cells. Nature, 509(7501), 487–491.

37. Jason, S. L., & Yusa, K. (2019). Genome-wide CRISPR-Cas9 screening in mammalian cells. Methods, 164, 29–35.

38. Bock, C., Datlinger, P., Chardon, F., Coelho, M.A., Dong, M.B., Lawson, K.A., Lu, T., Maroc, L., Norman, T.M., Song, B., & Stanley, G. (2022). High-content CRISPR screening. Nature Reviews Methods Primers, 2(1), 1–23.

39. Wang, T., Wei, J. J., Sabatini, D. M., & Lander, E. S. (2014). Genetic screens in human cells using the CRISPR-Cas9 system. Science, 343(6166), 80–84.

40. Terekhov, S.S., Eliseev, I.E., Ovchinnikova, L.A., Kabilov, M.R., Prjibelski, A.D., Tupikin, A.E., Smirnov, I.V., Belogurov Jr, A.A., Severinov, K.V., Lomakin, Y.A., & Altman, S. (2020). Liquid drop of DNA libraries reveals total genome information. Proceedings of the National Academy of Sciences, 117(44), 27300–27306.

41. Williams, R., Peisajovich, S. G., Miller, O. J., Magdassi, S., Tawfik, D. S., & Griffiths, A. D. (2006). Amplification of complex gene libraries by emulsion PCR. Nature methods, 3(7), 545–550.

42. Griffiths, A. D., & Tawfik, D. S. (2006). Miniaturising the laboratory in emulsion droplets. Trends in Biotechnology, 24(9), 395–402.

43. Shao, K., Ding, W., Wang, F., Li, H., Ma, D., & Wang, H. (2011). Emulsion PCR: a high efficient way of PCR amplification of random DNA libraries in aptamer selection. PLoS One, 6(9), e24910.

44. Paegel, B. M., & Joyce, G. F. (2010). Microfluidic compartmentalized directed evolution. Chemistry & Biology, 17, 717–724.

45. Hindson, B.J., Ness, K.D., Masquelier, D.A., Belgrader, P., Heredia, N.J., Makarewicz, A.J., Bright, I.J., Lucero, M.Y., Hiddessen, A.L., Legler, T.C., & Kitano, T.K. (2011). High-throughput droplet digital PCR system for absolute quantitation of DNA copy number. Analytical Chemistry, 83(22), 8604–8610.

46. Tawfik, D. S., & Griffiths, A. D. (1998). Man-made cell-like compartments for molecular evolution. Nature Biotechnology, 16(7), 652–656.

47. Bernath, K., Hai, M., Mastrobattista, E., Griffiths, A. D., Magdassi, S., & Tawfik, D. S. (2004). In vitro compartmentalization by double emulsions: sorting and gene enrichment by fluorescence activated cell sorting. Analytical Biochemistry, 325(1), 151–157.

48. Miller, O.J., Bernath, K., Agresti, J.J., Amitai, G., Kelly, B.T., Mastrobattista, E., Taly, V., Magdassi, S., Tawfik, D.S., & Griffiths, A.D. (2006). Directed evolution by in vitro compartmentalization. Nature Methods, 3(7), 561–570.

49. Schaerli, Y., & Hollfelder, F. (2009). The potential of microfluidic water-in-oil droplets in experimental biology. Molecular Biosystems, 5(12), 1392–1404.

50. Taly, V., Kelly, B. T., & Griffiths, A. D. (2007). Droplets as microreactors for high-throughput biology. ChemBioChem, 8(3), 263–272.

51. Kocak, D. D., Josephs, E. A., Bhandarkar, V., Adkar, S. S., Kwon, J. B., & Gersbach, C. A. (2019). Increasing the specificity of CRISPR systems with engineered RNA secondary structures. Nature Biotechnology, 37(6), 657–666.

52. Herring-Nicholas, A., Dimig, H., Roesing, M. R., & Josephs, E. A. (2024). Selection of extended CRISPR RNAs with enhanced targeting and specificity. Communications Biology, 7(1), 86.

53. Koonin, E. V., Gootenberg, J. S., & Abudayyeh, O. O. (2023). Discovery of diverse CRISPR-Cas systems and expansion of the genome engineering toolbox. Biochemistry, 62(24), 3465–3487.

54. Magoč, T., & Salzberg, S. L. (2011). FLASH: Fast length adjustment of short reads to improve genome assemblies. Bioinformatics, 27, 2957–2963.

55. Cock, P.J., Antao, T., Chang, J.T., Chapman, B.A., Cox, C.J., Dalke, A., Friedberg, I., Hamelryck, T., Kauff, F., Wilczynski, B., & De Hoon, M.J. (2009). Biopython: freely available Python tools for computational molecular biology and bioinformatics. Bioinformatics, 25(11), 1422.

56. Smith, T. F., & Waterman, M. S. (1981). Identification of common molecular subsequences. Journal of Molecular Biology, 147(1), 195–197.

