## Supporting Information for "Compartmentalized CRISPR Reactions (CCR) for High-Throughput Screening of Guide RNA Potency and Specificity"

**Materials and Methods**

*DNA oligonucleotides:*

DNA sequences for all oligonucleotides are listed in Supplementary Note 1-4. The targets/gRNAs and off-targets sequences are listed in Table S1 and Table S2.

**Table S1. 18 possible targets/gRNAs capable of targeting sequences within the 126 bp exon 1 of the human gene EMX1 (Chr2:72933825-72933951 in the hg38 assembly) as predicted by various CRISPR gRNA design algorithms [1-10]. Related to Figure 3.**

| Targets/gRNAs | Sequences |
| --- | --- |
| 1 | GCTTCGTGGCAATGCGCCACCGG |
| 2 | GCGCCACCGGTTGATGTGATGGG |
| 3 | GTGGCGCATTGCCACGAAGCAGG |
| 4 | TGCGCCACCGGTTGATGTGATGG |
| 5 | GCTCCCATCACATCAACCGGTGG |
| 6 | TTGCCACGAAGCAGGCCAATGGG |
| 7 | CTCCCCATTGGCCTGCTTCGTGG |
| 8 | GACATCGATGTCCTCCCCATTGG |
| 9 | TGCCACGAAGCAGGCCAATGGGG |
| 10 | AGGGCTCCCATCACATCAACCGG |
| 11 | CACGAAGCAGGCCAATGGGGAGG |
| 12 | TTCTTCTTCTGCTCGGACTCAGG |
| 13 | ATTGCCACGAAGCAGGCCAATGG |
| 14 | GGAGCCCTTCTTCTTCTGCTCGG |
| 15 | TGAGTCCGAGCAGAAGAAGAAGG |
| 16 | GAGTCCGAGCAGAAGAAGAAGGG |
| 17 | CGGCAGAAGCTGGAGGAGGAAGG |
| 18 | GGCAGAAGCTGGAGGAGGAAGGG |

**Table S2. 18 predicated off-target sites for four of the gRNAs from CRISPOR [11] as well as gRNA 16. Related to Figure 4.**

| # | Target/gRNAs | OffTarget |
| --- | --- | --- |
| 1 | 1 | GCTTTGTGGCAATGCACAACTGG |
| 2 | 1 | GCTTCGTGTCAATTTGCCACGGG |
| 3 | 2 | GGGCCATGGGTTGATGTGATGAG |
| 4 | 2 | GTGCCACCAGTTGATCTGATGGG |
| 5 | 3 | GTGGCTCATTGCCAAGGAGCCGG |
| 6 | 3 | GTGTGGCATTGACACGAAGCTGA |
| 7 | 3 | ATGGCACATTGCCACTAAGCAGG |
| 8 | 5 | GCTCCCATCACATCACCCTGAGG |
| 9 | 5 | TCTCCCATCACATCCACCAGTGG |
| 10 | 5 | GCTGCCCTCACATCAACAGGTGG |
| 11 | 5 | GATCCCATCACATCCACAGGTGG |
| 12 | 5 | GCTTCCATCACAGCAGCCGGGGG |
| 13 | 5 | GCCCCCATCAGATCTACCGGAGG |
| 14 | 5 | GCTCCCCTCACATGAACCTGAGG |
| 15 | 16 | GAGTTAGAGCAGAAGAAGAAAGG |
| 16 | 16 | GAGTCTAAGCAGAAGAAGAAGAG |
| 17 | 16 | GAGTCCTAGCAGGAGAAGAAGAG |
| 18 | 16 | GAGGCCGAGCAGAAGAAAGACGG |

***Protocol for Single Pot in vitro CRISPR-induced cleavage Reactions***

DNA oligonucleotides were purchased were purchased from Integrated DNA Technologies (IDT) and then resuspended to a stock concentration of 100 μM. DNA oligonucleotides were further diluted to a concentration of 300 fmol which is visible in agrarose gel stained with SYBR Safe dye. All the reagents were then mixed in the following order to set up CRISPR-mediated cleavage reaction: 7 μL nuclease-free water, 1 μL DNA Oligonucleotides (300 fmol), 2 μL NEB r 3.1 Buffer (10X), 1 μL RNAse Inhibitor (New England Biolabs #M0314S), 0.4 μL (2.5 mM) ribonucleotide triphosphate mix (NEB: N0466S), 3 μL (1000nM) Cas9 nuclease, *S. pyogenes* ( (NEB; M0386T) , 2 μL T7 RNA polymerase (New England BioLabs). The reaction mixture is then incubated for a duration of 3 hours at 37°C. This is followed by proteinase K digestion, where 1 μL of proteinase K (ThermoFisher kit #EO0491) is added, and the mixture is incubated at 56°C for 10 minutes. Subsequently, 2 μL of RNAse A is added to degrade the sgRNA molecules. The resulting products are then separated on a 3% agarose gel, stained with SYBR Safe, and analyzed using ImageJ.

***Protocol for Single Pot CRISPR-induced cleavage Reactions in Emulsion***

Water-oil emulsions were produced using the bulk mixing of water and oil phase following the procedure outlined by Williams et al.^41^ The oil phase was prepared by thoroughly mixing the following components, 2.25 ml -Span 80 4.5% (vol/vol), 200 µl -Tween 80 0.4% (vol/vol), 25 µl- Triton X-100 0.05% (vol/vol) and Mineral oil to 50 ml. A modification in the protocol was introduced for the aqueous phase, which comprised the following reagents for the CRISPR-induced cleavage reaction: 216.6 μL nuclease-free water, 1 μL DNA oligonucleotides (1 fmol/0.1 fmol), 26 μL NEB r 3.1 Buffer (10X), 5 μL RNAse Inhibitor (New England Biolabs #M0314S), 5.2 μL (2.5 mM) ribonucleotide triphosphate mix (NEB: N0466S), 1μL (1000nM) Cas9 nuclease, S. pyogenes (NEB; M0386T), 5.2 μL T7 RNA polymerase (New England BioLabs).

In a cryovial with a magnetic stir bar on a magnetic plate at an intermediate setting (1150 rpm), 400 μL of the oil phase was placed. Subsequently, 200 μL of the aqueous reaction mixture was slowly added to the oil phase over a 5-minute duration. It is important to note that the entire setup was maintained in a cold room at 4°C during the emulsification process. After this step, a stable water-oil emulsion was formed, followed by an 8-hour incubation at 37°C. The emulsion was then disrupted by centrifugation (15,000 rpm, 10 min) to recover the water phase, while the oil phase was pipetted out. The DNA library from the aqueous phase was extracted through a cleanup process with magnetic beads added in a 0.9X ratio to the aqueous phase. The purified DNA library was subsequently enriched using the NEBNext® Ultra™ II DNA Library Prep Kit for Illumina® (NEB# E7645S), followed by PCR with NEBNext® Multiplex Oligos for Illumina® (NEB# E7730S). The schematic of the process is depicted in Figure S1. The resulting products were also separated on a 3% agarose gel, stained with SYBR Safe, and then submitted for NGS sequencing.

***In vitro digestion reactions***

DNA targets containing the target sequences were synthesized by Twist Bioscience. Subsequently, they were PCR amplified using the provided universal primers, purified, and resuspended in nuclease-free water to to a concentration of 100 nM. For each reaction, three technical replicates were prepared in the following order: 7 μL of nuclease-free water, 1 μL of the target DNA substrate (100 nM), 1 μL of 10x Cas9 Nuclease Reaction buffer (composed of 200 mM HEPES, 1 M NaCl, 50 mM MgCl2, and 1 mM EDTA with a pH of 6.5 at 25°C), and 1 μL of Cas9-RNP (1 mM). The assembled reactions were then incubated for 1 hour at 37°C, followed by digestion with proteinase K (ThermoFisher enzyme #EO0491) (1 μL) at 56°C for 10 minutes. The resulting products were resolved on a 3% agarose gel stained with SYBR Gold and analyzed using ImageJ. For analysis of the gel image in Image J, the fluorescence intensity was normalized by length of the DNA fragments, and fraction cleaved was calculated using the equation:

$$Fraction cleaved= \left( \frac{Normalised cleaved band intensity}{\sum Normalised Cleaved and Uncleaved band Intensities} \right)\times100$$

***Dynamic light Scattering (DLS) Measurement***

The size of water droplets in water-oil emulsions were measured using a dynamic light scattering (DLS) instrument, Zetasizer Nano ZS90 (Malvern instruments, Malvern, UK). The average value of 3 run was taken as a final result. The stability of the water-oil emulsion was tested by measuring the water droplets immediately after emulsion formation and after a interval of 8 hours.

***Analysis of the NGS (Illumina) data and (Nanopore Sequencing) data***

The NGS sequencing data had sequences with pre-trimmed adaptors and barcode sequences. The paired end was then merged into longer, single reads using FLASH (Fast Length Adjustment of SHort reads) [12] bioinformatic tool. A script was written in Python that used Bio-python [13] package for reading the FASTQ files. Afterward, the sequences were divided into three categories: Uncut, Spacer Distal-cut (cleaved at the on-target site), and Spacer Proximal-cut (cleaved at the off-target site). Subsequently, top candidates in each category were refined based on alignment scores obtained through the Smith-Waterman algorithm [14] to the reference sequences. For the analysis of results from extended guideRNAs (x-gRNAs) screening experiment, a Python script was written to extract 8-nucleotide sequences (extended gRNA extensions) adjacent to the (T7 Promoter + off-target) sequence from the filtered large fragments list and stored them as a CSV file.

The reference code used for the analysis of the NGS data is provided at https://github.com/neelarka/CCR-Analysis

**Supplementary Figures**

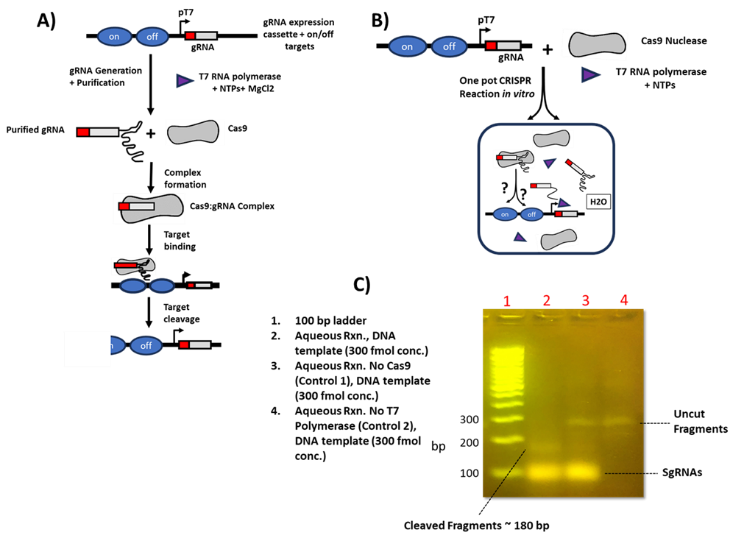

**Figure S1.** Single-pot CRISPR reaction conducted in an aqueous bulk phase, eliminating the necessity for external CRISPR gRNA synthesis. A) The standard protocol utilized for CRISPR- Cas9 cleavage assays. B) Our developed protocol enables performing the CRISPR- Cas9 cleavage assays in a one pot manner C) Representative agarose gel demonstrates the feasibility of one pot CRISPR- Cas9 cleavage assays. Samples without Cas9 effector and T7 polymerase are two controls here. DNA template concentration is ~ 349.2 femtomoles.

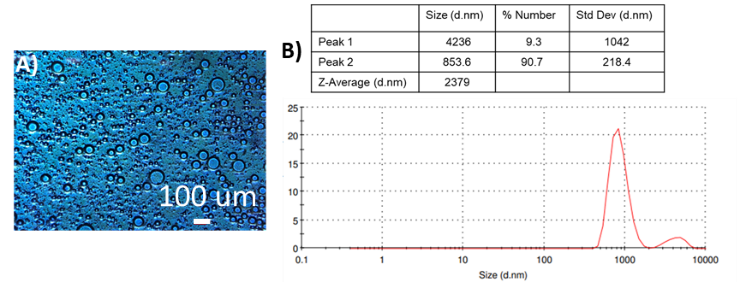

**Figure S2.** A) Microemulsions generated observed under optical microscope. B) Size of droplets in micro-emulsions measured through Dynamic Light Scattering (DLS*)*.

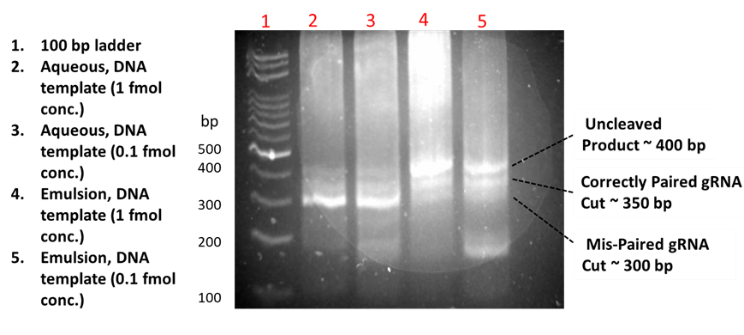

**Figure S3.** Validation of Compartmentalized CRISPR reactions (CCR) technique. Representative agarose gel image shows the results of our developed one pot CRISPR reactions performed in either under standard reaction conditions (with no emulsions) and within the emulsions. Lane details are as provided in the Figure. Note, each designed pair of DNA molecules contained a gRNA express cassette, its target on the distal end of the DNA molecule, and the target for the other paired gRNA positioned proximally between its target and the promotor.  If DNA molecules were isolated (as expected in micro-emulsions), we expect to see DNA fragments with cuts only on at the distal target sites (~350bp), while if both DNA pairs were mixed (as in standard in vitro condition), we expect DNA fragments cut at the proximal target site (~300 bp).

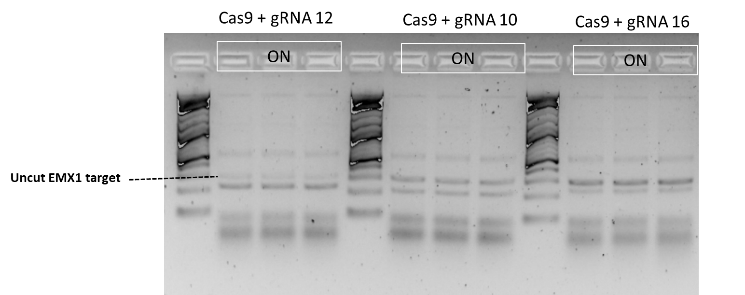

**Figure S4.** In vitro validation of the top three gRNA candidates capable of targeting sequences within the 126 bp exon 1 of the human gene EMX1 (Chr2:72933825-72933951 in the hg38 assembly) screened for on-target cleavage efficiency using Compartmentalized CRISPR reactions (CCR) approach. Note gRNA 12 has higher efficiency compared to gRNA 10 and gRNA 16. However, only gRNA 16 is only well-characterized in literature [15, 16] and confirmed to have high efficacy. The uncut target derived from EMX1 gene (by PCR) is around 300 bp and the cleaved DNA post in vitro CRISPR assay is around 200 bp. The additional band around 500bp corresponds to PCR byproducts derived from the EMX1 gene.

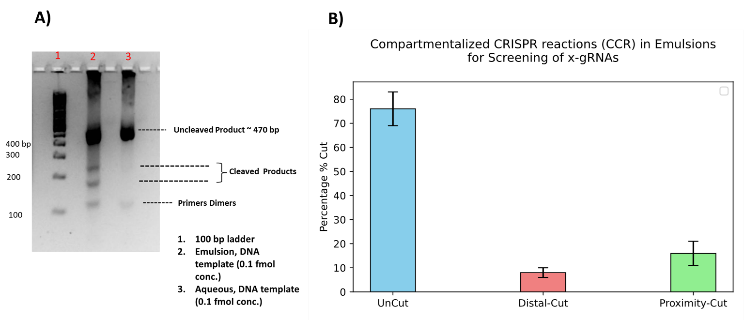

**Figure S5.** Compartmentalized CRISPR reactions (CCR) for screening of x-gRNAs. Designed DNA library comprised of all possible 8 nt’s combinations for 5’- extensions (65,536 possible sequences), for EMX1 gRNA 16 on a DNA molecule which had its target on the distal end and one each of the four most active off-target sites at the proximal site. A) Agarose gel image for the CRISPR cleavage reaction (one pot) in either under standard reaction conditions (with no emulsions) and within the emulsions with the DNA library designed for screening of x-gRNAs. Lane details are as provided in the figure. B) Analysis of the NGS result revealed that majority of the templates were uncut, while a smaller fraction, about 3590, efficient and specific extended gRNAs (x-gRNAs) were identified, which exhibited cleavage exclusively at the on-target sites and generated “Spacer Distal cut” DNA strands.

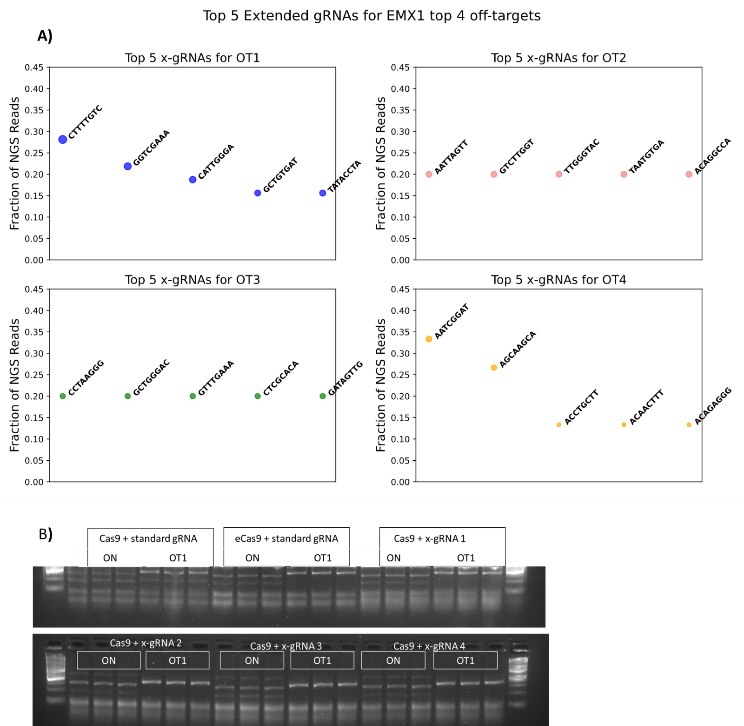

**Figure S6.** A) Determination of top “good” x-gRNAs from Next-Generation Sequencing (NGS). B) In vitro validation of the top four extended guide RNA (x-gRNAs) candidates (in triplicate) for EMX1 off-target OT1. Representative agarose gel shows RNP cleavage activity for EMX1 sgRNA with both Cas9 and Engineered Cas9 (eCas9) as well as screened x-gRNAs with Cas9. The uncut target is around 300 bp both for targets and off-targets and the cleaved DNA is around 200 bp.

**Supplementary Notes**

***Supplementary Note 1. EMX1 for in vitro validation related to Figure 3***

  5’-TATGTAGCCTCAGTCTTCCCATCAGGCTCTCAGCTCAGCCTGAGTGTTGA GGCCCCAGTGGCTGCTCTGGGGGCCTCCTGAGTTTCTCATCTGTGCCCCT CCCTCCCTGGCCCAGGTGAAGGTGTGGTTCCAGAACCGGAGGACAAAGTA CAAACGGCAGAAGCTGGAGGAGGAAGGGCCTGAGTCCGAGCAGAAGAAGA AGGGCTCCCATCACATCAACCGGTGGCGCATTGCCACGAAGCAGGCCAAT GGGGAGGACATCGATGTCACCTCCAATGACTAGGGTGGGCAACCACAAACC

***Supplementary Note 2.  DNA library related to Figure 2***

1. GACATCACCTCCCACAACGACGAAAAGAGGAGGAAGGGCCTGAGTCCGAGCAGAAGAAGAAGGGCTCCCATCACATCAATGGGCTTTGGAAAGGGGGTGGGGGGAGTTTGCTCCTGGACCCCCTATTTCTGATAATACGACTCACTATAGGAGTCCGAGCAGAAGAAGAAGTTTTAGAGCTAGAAATAGCAAGTTAAAATAAGGCTAGTCCGTTATCAACTTGAAAAAGTGGCACCGAGTCGGTGCTTTTTTAAACGAGGCGAGTTTACGGGTTGTTA
2. GACATCACCTCCCACAACGACGAGGGTGGGCTTTGGAAAGGGGGTGGGGGGAGTTTGCTCCTGGACCCCCTATTTCTGAGAGGAGGAAGGGCCTGAGTCCGAGCAGAAGAAGAAGGGCTCCCATCACATCAATAATACGACTCACTATAGGGGTGGGGGGAGTTTGCTCCGTTTTAGAGCTAGAAATAGCAAGTTAAAATAAGGCTAGTCCGTTATCAACTTGAAAAAGTGGCACCGAGTCGGTGCTTTTTTGGGCGAGGCGAGTTTACGGGTTGTTA

***Supplementary Note 3. On target DNA library related to Figure 3***

1) GACATCACCTCCCACAACGACGAAAACCTCCCCATTGGCCTGCTTCGTGGCAATGCGCCACCGGTTGATGTGATGGGAGGGATCTTATAGAGCCTGTGTGGACCCCCTATTTCTGATAATACGACTCACTATAGGCTTCGTGGCAATGCGCCACGTTTTAGAGCTAGAAATAGCAAGTTAAAATAAGGCTAGTCCGTTATCAACTTGAAAAAGTGGCACCGAGTCGGTGCTTTTTTAAACGAGGCGAGTTTACGGGTTGTTA
2) GACATCACCTCCCACAACGACGAAAACTGCTTCGTGGCAATGCGCCACCGGTTGATGTGATGGGAGCCCTTCTTCTTCTGGATCTTATAGAGCCTGTGTGGACCCCCTATTTCTGATAATACGACTCACTATAGGCGCCACCGGTTGATGTGATGTTTTAGAGCTAGAAATAGCAAGTTAAAATAAGGCTAGTCCGTTATCAACTTGAAAAAGTGGCACCGAGTCGGTGCTTTTTTAAACGAGGCGAGTTTACGGGTTGTTA
3) GACATCACCTCCCACAACGACGAAAACCATCACATCAACCGGTGGCGCATTGCCACGAAGCAGGCCAATGGGGAGGACAGGATCTTATAGAGCCTGTGTGGACCCCCTATTTCTGATAATACGACTCACTATAGGTGGCGCATTGCCACGAAGCGTTTTAGAGCTAGAAATAGCAAGTTAAAATAAGGCTAGTCCGTTATCAACTTGAAAAAGTGGCACCGAGTCGGTGCTTTTTTAAACGAGGCGAGTTTACGGGTTGTTA
4) GACATCACCTCCCACAACGACGAAAACCTGCTTCGTGGCAATGCGCCACCGGTTGATGTGATGGGAGCCCTTCTTCTTCGGATCTTATAGAGCCTGTGTGGACCCCCTATTTCTGATAATACGACTCACTATAGTGCGCCACCGGTTGATGTGAGTTTTAGAGCTAGAAATAGCAAGTTAAAATAAGGCTAGTCCGTTATCAACTTGAAAAAGTGGCACCGAGTCGGTGCTTTTTTAAACGAGGCGAGTTTACGGGTTGTTA
5) GACATCACCTCCCACAACGACGAAAAAGCAGAAGAAGAAGGGCTCCCATCACATCAACCGGTGGCGCATTGCCACGAAGGGATCTTATAGAGCCTGTGTGGACCCCCTATTTCTGATAATACGACTCACTATAGGCTCCCATCACATCAACCGGGTTTTAGAGCTAGAAATAGCAAGTTAAAATAAGGCTAGTCCGTTATCAACTTGAAAAAGTGGCACCGAGTCGGTGCTTTTTTAAACGAGGCGAGTTTACGGGTTGTTA
6) GACATCACCTCCCACAACGACGAAAATCAACCGGTGGCGCATTGCCACGAAGCAGGCCAATGGGGAGGACATCGATGTCGGATCTTATAGAGCCTGTGTGGACCCCCTATTTCTGATAATACGACTCACTATAGTTGCCACGAAGCAGGCCAATGTTTTAGAGCTAGAAATAGCAAGTTAAAATAAGGCTAGTCCGTTATCAACTTGAAAAAGTGGCACCGAGTCGGTGCTTTTTTAAACGAGGCGAGTTTACGGGTTGTTA
7) GACATCACCTCCCACAACGACGAAAAGGTGACATCGATGTCCTCCCCATTGGCCTGCTTCGTGGCAATGCGCCACCGGTGGATCTTATAGAGCCTGTGTGGACCCCCTATTTCTGATAATACGACTCACTATAGCTCCCCATTGGCCTGCTTCGGTTTTAGAGCTAGAAATAGCAAGTTAAAATAAGGCTAGTCCGTTATCAACTTGAAAAAGTGGCACCGAGTCGGTGCTTTTTTAAACGAGGCGAGTTTACGGGTTGTTA
8) GACATCACCTCCCACAACGACGAAAACTAGTCATTGGAGGTGACATCGATGTCCTCCCCATTGGCCTGCTTCGTGGCAAGGATCTTATAGAGCCTGTGTGGACCCCCTATTTCTGATAATACGACTCACTATAGGACATCGATGTCCTCCCCATGTTTTAGAGCTAGAAATAGCAAGTTAAAATAAGGCTAGTCCGTTATCAACTTGAAAAAGTGGCACCGAGTCGGTGCTTTTTTAAACGAGGCGAGTTTACGGGTTGTTA
9) GACATCACCTCCCACAACGACGAAAACAACCGGTGGCGCATTGCCACGAAGCAGGCCAATGGGGAGGACATCGATGTCAGGATCTTATAGAGCCTGTGTGGACCCCCTATTTCTGATAATACGACTCACTATAGTGCCACGAAGCAGGCCAATGGTTTTAGAGCTAGAAATAGCAAGTTAAAATAAGGCTAGTCCGTTATCAACTTGAAAAAGTGGCACCGAGTCGGTGCTTTTTTAAACGAGGCGAGTTTACGGGTTGTTA
10) GACATCACCTCCCACAACGACGAAAACCGAGCAGAAGAAGAAGGGCTCCCATCACATCAACCGGTGGCGCATTGCCACGGGATCTTATAGAGCCTGTGTGGACCCCCTATTTCTGATAATACGACTCACTATAGAGGGCTCCCATCACATCAACGTTTTAGAGCTAGAAATAGCAAGTTAAAATAAGGCTAGTCCGTTATCAACTTGAAAAAGTGGCACCGAGTCGGTGCTTTTTTAAACGAGGCGAGTTTACGGGTTGTTA
11) GACATCACCTCCCACAACGACGAAAACCGGTGGCGCATTGCCACGAAGCAGGCCAATGGGGAGGACATCGATGTCACCTGGATCTTATAGAGCCTGTGTGGACCCCCTATTTCTGATAATACGACTCACTATAGCACGAAGCAGGCCAATGGGGGTTTTAGAGCTAGAAATAGCAAGTTAAAATAAGGCTAGTCCGTTATCAACTTGAAAAAGTGGCACCGAGTCGGTGCTTTTTTAAACGAGGCGAGTTTACGGGTTGTTA
12) GACATCACCTCCCACAACGACGAAAAATGTGATGGGAGCCCTTCTTCTTCTGCTCGGACTCAGGCCCTTCCTCCTCCAGGGATCTTATAGAGCCTGTGTGGACCCCCTATTTCTGATAATACGACTCACTATAGTTCTTCTTCTGCTCGGACTCGTTTTAGAGCTAGAAATAGCAAGTTAAAATAAGGCTAGTCCGTTATCAACTTGAAAAAGTGGCACCGAGTCGGTGCTTTTTTAAACGAGGCGAGTTTACGGGTTGTTA
13) GACATCACCTCCCACAACGACGAAAAATCAACCGGTGGCGCATTGCCACGAAGCAGGCCAATGGGGAGGACATCGATGTGGATCTTATAGAGCCTGTGTGGACCCCCTATTTCTGATAATACGACTCACTATAGATTGCCACGAAGCAGGCCAAGTTTTAGAGCTAGAAATAGCAAGTTAAAATAAGGCTAGTCCGTTATCAACTTGAAAAAGTGGCACCGAGTCGGTGCTTTTTTAAACGAGGCGAGTTTACGGGTTGTTA
14) GACATCACCTCCCACAACGACGAAAACCGGTTGATGTGATGGGAGCCCTTCTTCTTCTGCTCGGACTCAGGCCCTTCCTGGATCTTATAGAGCCTGTGTGGACCCCCTATTTCTGATAATACGACTCACTATAGGGAGCCCTTCTTCTTCTGCTGTTTTAGAGCTAGAAATAGCAAGTTAAAATAAGGCTAGTCCGTTATCAACTTGAAAAAGTGGCACCGAGTCGGTGCTTTTTTAAACGAGGCGAGTTTACGGGTTGTTA
15) GACATCACCTCCCACAACGACGAAAAGGAGGAGGAAGGGCCTGAGTCCGAGCAGAAGAAGAAGGGCTCCCATCACATCAGGATCTTATAGAGCCTGTGTGGACCCCCTATTTCTGATAATACGACTCACTATAGTGAGTCCGAGCAGAAGAAGAGTTTTAGAGCTAGAAATAGCAAGTTAAAATAAGGCTAGTCCGTTATCAACTTGAAAAAGTGGCACCGAGTCGGTGCTTTTTTAAACGAGGCGAGTTTACGGGTTGTTA
16) GACATCACCTCCCACAACGACGAAAAGAGGAGGAAGGGCCTGAGTCCGAGCAGAAGAAGAAGGGCTCCCATCACATCAAGGATCTTATAGAGCCTGTGTGGACCCCCTATTTCTGATAATACGACTCACTATAGGAGTCCGAGCAGAAGAAGAAGTTTTAGAGCTAGAAATAGCAAGTTAAAATAAGGCTAGTCCGTTATCAACTTGAAAAAGTGGCACCGAGTCGGTGCTTTTTTAAACGAGGCGAGTTTACGGGTTGTTA
17) GACATCACCTCCCACAACGACGAAAAAGGACAAAGTACAAACGGCAGAAGCTGGAGGAGGAAGGGCCTGAGTCCGAGCAGGATCTTATAGAGCCTGTGTGGACCCCCTATTTCTGATAATACGACTCACTATAGCGGCAGAAGCTGGAGGAGGAGTTTTAGAGCTAGAAATAGCAAGTTAAAATAAGGCTAGTCCGTTATCAACTTGAAAAAGTGGCACCGAGTCGGTGCTTTTTTAAACGAGGCGAGTTTACGGGTTGTTA
18) GACATCACCTCCCACAACGACGAAAAGGACAAAGTACAAACGGCAGAAGCTGGAGGAGGAAGGGCCTGAGTCCGAGCAGGGATCTTATAGAGCCTGTGTGGACCCCCTATTTCTGATAATACGACTCACTATAGGGCAGAAGCTGGAGGAGGAAGTTTTAGAGCTAGAAATAGCAAGTTAAAATAAGGCTAGTCCGTTATCAACTTGAAAAAGTGGCACCGAGTCGGTGCTTTTTTAAACGAGGCGAGTTTACGGGTTGTTA

***Supplementary Note 4. On-OFF DNA library related to Figure 4***

****************ON***********************************                           ****************OFF**********************************

1. GACATCACCTCCCACAACGACGAAAACCTCCCCATTGGCCTGCTTCGTGGCAATGCGCCACCGGTTGATGTGATGGGAGTGGGCTTTGGAAAGGCTTATGGCATGGGATATGTCTGGATGTGCTTTGTGGCAATGCACAACTGGTCAGTGGGAAGGGACATCTTATAGAGCCTGTGTGGACCCCCTATTTCTGATAATACGACTCACTATAGGCTTCGTGGCAATGCGCCACGTTTTAGAGCTAGAAATAGCAAGTTAAAATAAGGCTAGTCCGTTATCAACTTGAAAAAGTGGCACCGAGTCGGTGCTTTTTTAAACGAGGCGAGTTTACGGGTTGTTA
2. GACATCACCTCCCACAACGACGAAAACCTCCCCATTGGCCTGCTTCGTGGCAATGCGCCACCGGTTGATGTGATGGGAGTGGGCTTTGGAAAGGCTTATGGCATGGCCTTAACTCGAGGTGCCCGTGGCAAATTGACACGAAGCTGTCCACCTGTTGGAATCTTATAGAGCCTGTGTGGACCCCCTATTTCTGATAATACGACTCACTATAGGCTTCGTGGCAATGCGCCACGTTTTAGAGCTAGAAATAGCAAGTTAAAATAAGGCTAGTCCGTTATCAACTTGAAAAAGTGGCACCGAGTCGGTGCTTTTTTAAACGAGGCGAGTTTACGGGTTGTTA
3. GACATCACCTCCCACAACGACGAAAACTGCTTCGTGGCAATGCGCCACCGGTTGATGTGATGGGAGCCCTTCTTCTTCTTGGGCTTTGGAAAGGCTTATGGCATGGCTCTAGAGACACATGCTCATCACATCAACCCATGGCCCAGAAACCTCCAGTGGATCTTATAGAGCCTGTGTGGACCCCCTATTTCTGATAATACGACTCACTATAGGCGCCACCGGTTGATGTGATGTTTTAGAGCTAGAAATAGCAAGTTAAAATAAGGCTAGTCCGTTATCAACTTGAAAAAGTGGCACCGAGTCGGTGCTTTTTTAAACGAGGCGAGTTTACGGGTTGTTA
4. GACATCACCTCCCACAACGACGAAAACTGCTTCGTGGCAATGCGCCACCGGTTGATGTGATGGGAGCCCTTCTTCTTCTTGGGCTTTGGAAAGGCTTATGGCATGGGAATCAACCTAGATGCCCATCAGATCAACTGGTGGCACACATACACTATGAAAATCTTATAGAGCCTGTGTGGACCCCCTATTTCTGATAATACGACTCACTATAGGCGCCACCGGTTGATGTGATGTTTTAGAGCTAGAAATAGCAAGTTAAAATAAGGCTAGTCCGTTATCAACTTGAAAAAGTGGCACCGAGTCGGTGCTTTTTTAAACGAGGCGAGTTTACGGGTTGTTA
5. GACATCACCTCCCACAACGACGAAAACCATCACATCAACCGGTGGCGCATTGCCACGAAGCAGGCCAATGGGGAGGACATGGGCTTTGGAAAGGCTTATGGCATGGAGCCGCAGCCTGGCCCCGGCTCCTTGGCAATGAGCCACCTCCTACCTGTCGCCATCTTATAGAGCCTGTGTGGACCCCCTATTTCTGATAATACGACTCACTATAGGTGGCGCATTGCCACGAAGCGTTTTAGAGCTAGAAATAGCAAGTTAAAATAAGGCTAGTCCGTTATCAACTTGAAAAAGTGGCACCGAGTCGGTGCTTTTTTAAACGAGGCGAGTTTACGGGTTGTTA
6. GACATCACCTCCCACAACGACGAAAACCATCACATCAACCGGTGGCGCATTGCCACGAAGCAGGCCAATGGGGAGGACATGGGCTTTGGAAAGGCTTATGGCATGGCAGTGTGCACCTCACTCAGCTTCGTGTCAATGCCACACCATGTGGGGAGTGTTATCTTATAGAGCCTGTGTGGACCCCCTATTTCTGATAATACGACTCACTATAGGTGGCGCATTGCCACGAAGCGTTTTAGAGCTAGAAATAGCAAGTTAAAATAAGGCTAGTCCGTTATCAACTTGAAAAAGTGGCACCGAGTCGGTGCTTTTTTAAACGAGGCGAGTTTACGGGTTGTTA
7. GACATCACCTCCCACAACGACGAAAACCATCACATCAACCGGTGGCGCATTGCCACGAAGCAGGCCAATGGGGAGGACATGGGCTTTGGAAAGGCTTATGGCATGGAAATGTTCTAGGACCATGGCACATTGCCACTAAGCAGGAGTCAAAAGACTACAATCTTATAGAGCCTGTGTGGACCCCCTATTTCTGATAATACGACTCACTATAGGTGGCGCATTGCCACGAAGCGTTTTAGAGCTAGAAATAGCAAGTTAAAATAAGGCTAGTCCGTTATCAACTTGAAAAAGTGGCACCGAGTCGGTGCTTTTTTAAACGAGGCGAGTTTACGGGTTGTTA
8. GACATCACCTCCCACAACGACGAAAAAGCAGAAGAAGAAGGGCTCCCATCACATCAACCGGTGGCGCATTGCCACGAAGTGGGCTTTGGAAAGGCTTATGGCATGGGAGGTGTCTCTGAAGCCTCAGGGTGATGTGATGGGAGCCCTGGTGCCCTCACTATCTTATAGAGCCTGTGTGGACCCCCTATTTCTGATAATACGACTCACTATAGGCTCCCATCACATCAACCGGGTTTTAGAGCTAGAAATAGCAAGTTAAAATAAGGCTAGTCCGTTATCAACTTGAAAAAGTGGCACCGAGTCGGTGCTTTTTTAAACGAGGCGAGTTTACGGGTTGTTA
9. GACATCACCTCCCACAACGACGAAAAAGCAGAAGAAGAAGGGCTCCCATCACATCAACCGGTGGCGCATTGCCACGAAGTGGGCTTTGGAAAGGCTTATGGCATGGGCCTGGCCAAGAAGTTCTCCCATCACATCCACCAGTGGATTCCCCTGCTGGGTATCTTATAGAGCCTGTGTGGACCCCCTATTTCTGATAATACGACTCACTATAGGCTCCCATCACATCAACCGGGTTTTAGAGCTAGAAATAGCAAGTTAAAATAAGGCTAGTCCGTTATCAACTTGAAAAAGTGGCACCGAGTCGGTGCTTTTTTAAACGAGGCGAGTTTACGGGTTGTTA
10. GACATCACCTCCCACAACGACGAAAAAGCAGAAGAAGAAGGGCTCCCATCACATCAACCGGTGGCGCATTGCCACGAAGTGGGCTTTGGAAAGGCTTATGGCATGGTCCCAGTAGGTGCATGCTGCCCTCACATCAACAGGTGGAGTCTATTTCCTCTCATCTTATAGAGCCTGTGTGGACCCCCTATTTCTGATAATACGACTCACTATAGGCTCCCATCACATCAACCGGGTTTTAGAGCTAGAAATAGCAAGTTAAAATAAGGCTAGTCCGTTATCAACTTGAAAAAGTGGCACCGAGTCGGTGCTTTTTTAAACGAGGCGAGTTTACGGGTTGTTA
11. GACATCACCTCCCACAACGACGAAAAAGCAGAAGAAGAAGGGCTCCCATCACATCAACCGGTGGCGCATTGCCACGAAGTGGGCTTTGGAAAGGCTTATGGCATGGGGTGACAAGTGGCAGGATCCCATCACATCCACAGGTGGCCTCGACTCTGGTCTATCTTATAGAGCCTGTGTGGACCCCCTATTTCTGATAATACGACTCACTATAGGCTCCCATCACATCAACCGGGTTTTAGAGCTAGAAATAGCAAGTTAAAATAAGGCTAGTCCGTTATCAACTTGAAAAAGTGGCACCGAGTCGGTGCTTTTTTAAACGAGGCGAGTTTACGGGTTGTTA
12. GACATCACCTCCCACAACGACGAAAAAGCAGAAGAAGAAGGGCTCCCATCACATCAACCGGTGGCGCATTGCCACGAAGTGGGCTTTGGAAAGGCTTATGGCATGGTCCCTTGTCCTGATGGCTTCCATCACAGCAGCCGGGGGCAACTCAGCCTGGACATCTTATAGAGCCTGTGTGGACCCCCTATTTCTGATAATACGACTCACTATAGGCTCCCATCACATCAACCGGGTTTTAGAGCTAGAAATAGCAAGTTAAAATAAGGCTAGTCCGTTATCAACTTGAAAAAGTGGCACCGAGTCGGTGCTTTTTTAAACGAGGCGAGTTTACGGGTTGTTA
13. GACATCACCTCCCACAACGACGAAAAAGCAGAAGAAGAAGGGCTCCCATCACATCAACCGGTGGCGCATTGCCACGAAGTGGGCTTTGGAAAGGCTTATGGCATGGATTGTTCAGTGATCTGCCCCCATCAGATCTACCGGAGGCCCTTGCCTCAACCTATCTTATAGAGCCTGTGTGGACCCCCTATTTCTGATAATACGACTCACTATAGGCTCCCATCACATCAACCGGGTTTTAGAGCTAGAAATAGCAAGTTAAAATAAGGCTAGTCCGTTATCAACTTGAAAAAGTGGCACCGAGTCGGTGCTTTTTTAAACGAGGCGAGTTTACGGGTTGTTA
14. GACATCACCTCCCACAACGACGAAAAAGCAGAAGAAGAAGGGCTCCCATCACATCAACCGGTGGCGCATTGCCACGAAGTGGGCTTTGGAAAGGCTTATGGCATGGGCTCCCTTCTACCCAGCTCCCCTCACATGAACCTGAGGGCCCTGTCAAGGTGGATCTTATAGAGCCTGTGTGGACCCCCTATTTCTGATAATACGACTCACTATAGGCTCCCATCACATCAACCGGGTTTTAGAGCTAGAAATAGCAAGTTAAAATAAGGCTAGTCCGTTATCAACTTGAAAAAGTGGCACCGAGTCGGTGCTTTTTTAAACGAGGCGAGTTTACGGGTTGTTA
15. GACATCACCTCCCACAACGACGAAAAGAGGAGGAAGGGCCTGAGTCCGAGCAGAAGAAGAAGGGCTCCCATCACATCAATGGGCTTTGGAAAGGCTTATGGCATGGTGCCTTTACTCCATGCCTTTCTTCTTCTGCTCTAACTCTGACAATCTGTCTTGATCTTATAGAGCCTGTGTGGACCCCCTATTTCTGATAATACGACTCACTATAGGAGTCCGAGCAGAAGAAGAAGTTTTAGAGCTAGAAATAGCAAGTTAAAATAAGGCTAGTCCGTTATCAACTTGAAAAAGTGGCACCGAGTCGGTGCTTTTTTAAACGAGGCGAGTTTACGGGTTGTTA
16. GACATCACCTCCCACAACGACGAAAAGAGGAGGAAGGGCCTGAGTCCGAGCAGAAGAAGAAGGGCTCCCATCACATCAATGGGCTTTGGAAAGGCTTATGGCATGGATTCATAGTAGACAAGAGTCTAAGCAGAAGAAGAAGAGAGCCACTACCCAACCATCTTATAGAGCCTGTGTGGACCCCCTATTTCTGATAATACGACTCACTATAGGAGTCCGAGCAGAAGAAGAAGTTTTAGAGCTAGAAATAGCAAGTTAAAATAAGGCTAGTCCGTTATCAACTTGAAAAAGTGGCACCGAGTCGGTGCTTTTTTAAACGAGGCGAGTTTACGGGTTGTTA
17. GACATCACCTCCCACAACGACGAAAAGAGGAGGAAGGGCCTGAGTCCGAGCAGAAGAAGAAGGGCTCCCATCACATCAATGGGCTTTGGAAAGGCTTATGGCATGGGGGCCAGCATGACCTGAGTCCTAGCAGGAGAAGAAGAGGCAGCCTAGAGTCTTATCTTATAGAGCCTGTGTGGACCCCCTATTTCTGATAATACGACTCACTATAGGAGTCCGAGCAGAAGAAGAAGTTTTAGAGCTAGAAATAGCAAGTTAAAATAAGGCTAGTCCGTTATCAACTTGAAAAAGTGGCACCGAGTCGGTGCTTTTTTAAACGAGGCGAGTTTACGGGTTGTTA
18. GACATCACCTCCCACAACGACGAAAAGAGGAGGAAGGGCCTGAGTCCGAGCAGAAGAAGAAGGGCTCCCATCACATCAATGGGCTTTGGAAAGGCTTATGGCATGGTCTTCTGCAAATGAGGAGGCCGAGCAGAAGAAAGACGGCGACAGATGTTGGGGATCTTATAGAGCCTGTGTGGACCCCCTATTTCTGATAATACGACTCACTATAGGAGTCCGAGCAGAAGAAGAAGTTTTAGAGCTAGAAATAGCAAGTTAAAATAAGGCTAGTCCGTTATCAACTTGAAAAAGTGGCACCGAGTCGGTGCTTTTTTAAACGAGGCGAGTTTACGGGTTGTTA

                          ****************ON***********************************                           ****************OFF**********************************

***Supplementary Note 4. Extended gRNAs (x-gRNAs) Library related to Figure 5***

1. GACATCACCTCCCACAACGACGAAAAGAGGAGGAAGGGCCTGAGTCCGAGCAGAAGAAGAAGGGCTCCCATCACATCAATGGGCTTTGGAAAGGcttatggcatggcaagacagattgtcaGAGTTAGAGCAGAAGAAGAAAGGcatggagtaaaggcaatcttatagagcctgtgTGGACCCCCTATTTCTGATAATACGACTCACTATAGNNNNNNNNGAGTCCGAGCAGAAGAAGAAGTTTTAGAGCTAGAAATAGCAAGTTAAAATAAGGCTAGTCCGTTATCAACTTGAAAAAGTGGCACCGAGTCGGTGCTTTTTTAAACGAGGCGAGTTTACGGGTTGTTA
2. GACATCACCTCCCACAACGACGAAAAGAGGAGGAAGGGCCTGAGTCCGAGCAGAAGAAGAAGGGCTCCCATCACATCAATGGGCTTTGGAAAGGcttatggcatggattcatagtagacaaGAGTCTAAGCAGAAGAAGAAGAGagccactacccaaccatctaagagagactgtgTGGACCCCCTATTTCTGATAATACGACTCACTATAGNNNNNNNNGAGTCCGAGCAGAAGAAGAAGTTTTAGAGCTAGAAATAGCAAGTTAAAATAAGGCTAGTCCGTTATCAACTTGAAAAAGTGGCACCGAGTCGGTGCTTTTTTAAACGAGGCGAGTTTACGGGTTGTTA
3. GACATCACCTCCCACAACGACGAAAAGAGGAGGAAGGGCCTGAGTCCGAGCAGAAGAAGAAGGGCTCCCATCACATCAATGGGCTTTGGAAAGGcttatggcatggtcttctgcaaatgagGAGGCCGAGCAGAAGAAAGACGGcgacagatgttggggggaggcaggtagctgtgTGGACCCCCTATTTCTGATAATACGACTCACTATAGNNNNNNNNGAGTCCGAGCAGAAGAAGAAGTTTTAGAGCTAGAAATAGCAAGTTAAAATAAGGCTAGTCCGTTATCAACTTGAAAAAGTGGCACCGAGTCGGTGCTTTTTTAAACGAGGCGAGTTTACGGGTTGTTA
4. GACATCACCTCCCACAACGACGAAAAGAGGAGGAAGGGCCTGAGTCCGAGCAGAAGAAGAAGGGCTCCCATCACATCAATGGGCTTTGGAAAGGcttatggcatgggggccagcatgacctGAGTCCTAGCAGGAGAAGAAGAGgcagcctagagtcttatcttatagagcctgtgTGGACCCCCTATTTCTGATAATACGACTCACTATAGNNNNNNNNGAGTCCGAGCAGAAGAAGAAGTTTTAGAGCTAGAAATAGCAAGTTAAAATAAGGCTAGTCCGTTATCAACTTGAAAAAGTGGCACCGAGTCGGTGCTTTTTTAAACGAGGCGAGTTTACGGGTTGTTA

**Supplementary References**

1. Doench, J. G., Fusi, N., Sullender, M., Hegde, M., Vaimberg, E. W., Donovan, K. F., Root, D. E. (2016). Optimized sgRNA design to maximize activity and minimize off-target effects of CRISPR-Cas9. Nature Biotechnology, 34, 184-191.
2. Moreno-Mateos, M., Vejnar, C., Beaudoin, J. D., et al. (2015). CRISPRscan: designing highly efficient sgRNAs for CRISPR-Cas9 targeting in vivo. Nature Methods, 12, 982–988.
3. Haeussler, M., Schönig, K., Eckert, H., et al. (2016). Evaluation of off-target and on-target scoring algorithms and integration into the guide RNA selection tool CRISPOR. Genome Biology, 17, 148.
4. Chuai, G., Ma, H., Yan, J., Chen, M., Hong, N., Xue, D., & Liu, Q. (2018). DeepCRISPR: optimized CRISPR guide RNA design by deep learning. Genome Biology, 19, 1-18.
5. Xu, H., Xiao, T., Chen, C. H., Li, W., Meyer, C. A., Wu, Q., & Liu, X. S. (2015). Sequence determinants of improved CRISPR sgRNA design. Genome Research, 25(8), 1147-1157.
6. Wong, N., Liu, W., & Wang, X. (2015). WU-CRISPR: characteristics of functional guide RNAs for the CRISPR/Cas9 system. Genome Biology, 16, 1-8.
7. Doench, J. G., Hartenian, E., Graham, D. B., Tothova, Z., Hegde, M., Smith, I., Sullender, M., et al. (2014). Rational design of highly active sgRNAs for CRISPR-Cas9–mediated gene inactivation. Nature Biotechnology, 32(12), 1262-1267.
8. Wang, D., Zhang, C., Wang, B., Li, B., Wang, Q., Liu, D., Wang, H., et al. (2019). Optimized CRISPR guide RNA design for two high-fidelity Cas9 variants by deep learning. Nature Communications, 10(1), 4284.
9. Fusi, N., Smith, I., Doench, J., & Listgarten, J. (2015). In silico predictive modeling of CRISPR/Cas9 guide efficiency. BioRxiv, 021568.
10. Chari, R., Mali, P., Moosburner, M., et al. (2015). Unraveling CRISPR-Cas9 genome engineering parameters via a library-on-library approach. Nature Methods, 12, 823–826.
11. Evaluation of off-target and on-target scoring algorithms and integration into the guide RNA selection tool CRISPOR. Genome biology, 17, 1-12. (2016). Evaluation of off-target and on-target scoring algorithms and integration into the guide RNA selection tool CRISPOR. Genome biology, 17, 1-12.
12. Magoč, T., & Salzberg, S. L. (2011). FLASH: Fast length adjustment of short reads to improve genome assemblies. Bioinformatics, 27, 2957-2963.
13. Cock, P.J., Antao, T., Chang, J.T., Chapman, B.A., Cox, C.J., Dalke, A., Friedberg, I., Hamelryck, T., Kauff, F., Wilczynski, B., & De Hoon, M.J. (2009). Biopython: freely available Python tools for computational molecular biology and bioinformatics. Bioinformatics, 25(11), 1422.
14. Smith, T. F., & Waterman, M. S. (1981). Identification of common molecular subsequences. Journal of Molecular Biology, 147(1), 195-197.
15. Kocak, D. D., Josephs, E. A., Bhandarkar, V., Adkar, S. S., Kwon, J. B., & Gersbach, C. A. (2019). Increasing the specificity of CRISPR systems with engineered RNA secondary structures. Nature Biotechnology, 37(6), 657-666.
16. Herring-Nicholas, A., Dimig, H., Roesing, M. R., & Josephs, E. A. (2024). Selection of extended CRISPR RNAs with enhanced targeting and specificity. Communications Biology, 7(1), 86.
